# Healthcare related activity following kidney transplantation: an observational cohort study

**DOI:** 10.1101/524421

## BACKGROUND

Renal transplantation is regarded as the gold standard of renal replacement therapy (RRT) with evidence of improved life expectancy, quality of life and superior cost-effectiveness when compared to dialysis (1–6). In recent years there has been a steady rise in the incidence of renal transplantation within most developed healthcare economies (7). Such has been the observed benefit that transplantation is now being considered across an increasingly wide range of patient age, BMI and comorbidities (8,9). With increased demand comes increased pressure on supply of deceased donor kidneys, for which the increased utilisation of extended criteria donation has been a positive step.

Despite superiority over dialysis, renal transplantation nonetheless involves invasive procedures, hospitalisation, frequent clinic attendances, heightened risk of intercurrent illnesses, including potentially life threatening infections, malignancy, or peri-operative mortality which may occur as a direct result of either the kidney transplant procedure or immunosuppressive therapy used to facilitate transplantation. Whilst data on patient and graft survival are routinely available, and rates of individual transplant-associated pathologies such as rejection, cytomegalovirus infection, or new onset diabetes are well described in the published literature, there is a paucity of data detailing what patients and services may expect in terms of all the combined downstream healthcare associated activities in the post-transplant period.

This is of particular importance when considering that transplantation has evolved considerably over the last decade, particularly in terms of the characteristics of both donors and recipients accepted for transplantation, so that much of the evidence on which services are based and quality is judged, is potentially outdated. Similarly, the future of renal transplantation is likely to involve further expansion in transplant numbers, with more complex donor and recipient combinations. These include recipients of increasingly higher body mass index, and kidneys from more ‘marginal’ deceased donors, such as the elderly, or higher perceived risk of infection (10–12). An understanding of the breadth and depth of healthcare related activity accrued on the patient’s journey following transplantation, and the associated clinical resources required to support it, is therefore advantageous to the individual and to the service provider, in terms of expectations, quality of care, and resource management.

In this study we describe a one-year cohort of transplanted patients followed up for 2 years post-transplant; aiming to capture the cumulative clinical activity and outcomes which impact on both patients and services alike. Further consideration is given to subgroups in which there are perceived to exist an excess of healthcare-associated activity.

## METHODS

All consecutive patients in the Glasgow Renal and Transplant Unit (UK) in receipt of a kidney transplant between 1^st^ of January and 31^st^ of December 2015 were studied. Patient demographics and events of interest (hospitalisation, outpatient clinics, imaging, biopsy procedures, and infections) were collated from electronic renal patient records for a period of up to 730 days (2 years) after transplantation. The date for each event was converted into a numerical day-post-transplant, allowing comparative visual representation of the activity.

The Glasgow Renal and Transplant Unit provide transplantation services for half of the Scottish population (NHS Greater Glasgow & Clyde, NHS Lanarkshire, NHS Dumfries & Galloway, and NHS Forth Valley Health Boards). Electronic health records capture data pertaining to every clinical encounter and healthcare-related activity undertaken in the West of Scotland, including all microbiology, virology, radiology, pathology, biochemistry and haematology tests; ensuring complete data capture.

Extended-criteria donors (ECDs) were defined as kidney donors aged ≥60 years, or donors who were aged 50 to 59 years and had two of the following three features: Hypertension, terminal serum creatinine >1.5 mg/dl (133mmol/L), or death from cerebrovascular accident (13). Standard-criteria donors (SCD) were defined as any deceased donor not encompassed in the above criteria.

The standard immunosuppression regimen comprised prednisolone 20mg daily (tapering to 5mg at by 3 months), mycophenolate mofetil (target 1000mg twice daily), Tacrolimus (Prograf) twice daily with target trough 5-8ng/ml, and Basiliximab (Simulect) 20mg day 1 and 4. In addition, co-trimoxazole 480mg daily as Pneumocystis jirovecci prophylaxis, and valganciclovir as prophylaxis against CMV in those at highest risk (donor CMV serology positive, recipient negative) was used for 6 months following transplantation (or treatment of acute rejection). Following discharge from hospital after transplantation, care was co-ordinated through the out-patient Acute Transplant Clinic run jointly by clinical nephrologists and transplant surgeons for the first year, after which care was delivered by their local nephrology team. All data were censored at the point of graft failure and return to dialysis or death.

Infection data included blood and urine cultures, clinically significant episodes of cytomegalovirus, BK virus, Pneumocystis jirovecci pneumonia, and Epstein Barr virus. Samples for blood cultures were incubated and monitored using BacTAlert (Biomerieux) and those which flagged positive were processed according to the Public Health England Standards for Microbiology Investigations (SMI) (14). Bacteraemic events were defined as positive microbiological growth of a pathogenic organism from blood cultures more than 14 days from any previous positive blood culture. Urinary tract infections (UTI) were diagnosed on urine culture with greater than 100,000 cfu/ml, more than 7 days from a prior UTI. Pneumocystis jirovecci pneumonia diagnosis required PCR detection of the organism with recorded antibiotic treatment. Viraemia with cytomegalovirus (CMV) and BK virus were deemed clinically significant if titres were quantified (rather than just “detected”), and greater than 100 copies/ml, respectively.

Measurements of white blood cell count (WCC) were considered to be as per the most recent blood sample for that individual patient, with all West of Scotland haematology results available; an average across all patients was then generated for each of the 730 days of the study period, for consideration alongside timing of infection episodes,

Protocol renal transplant biopsies were not performed in any patients. Transplant biopsy was performed in patients with significant deterioration in transplant function not explained by intercurrent illness, medicine nephrotoxicity, or abnormalities on ultrasound scan. Biopsies at the time of implantation were not included as they were felt to represent a component of the surgical procedure rather than subsequent healthcare activity.

Post-transplantation diabetes mellitus (PTDM) status was based on diagnoses entered onto the electronic patient record. PTDM is diagnosed in our unit on the basis of an oral glucose tolerance test, fasting laboratory glucose measure, or persisting requirement for insulin or other hypoglycaemic agents.

Data analysis was conducted with continuous variables described using means and standard deviations, or if not normally distributed, by medians and the interquartile range. Statistical analysis of categorical variables used the chi-square testing, whilst continuous variables were subjected to either Student’s t-test with a two-tailed hypothesis and significance level set at p<0.05 if normally distributed, or the Mann Whitney test if not normally distributed. The project was registered with NHS Greater Glasgow and Clyde as a clinical effectiveness project; due to the observational nature of this clinical service-related study, ethical approval was waived.

## RESULTS

### Demographics

132 consecutive patients underwent renal transplantation during 2015. Demographic data are presented in Table 1. Complete follow up for all recipients was available for the full 2 year study period. A total of 33/132 (25.0%) patients had a BMI >30kg/m^2^; this group were more likely to receive an ECD kidney 15/33 (45.5%), compared to 23/99 (23.2%) with BMI <30kg/m^2^ (OR 2.7 (95% CI 1.2, 6.3) p=0.03). Diabetes as the primary renal diagnosis (the documented cause of ESRD) demonstrated increasing prevalence with recipient BMI category, accounting for 4/58 (6.9%) patients with BMI <25kg/m^2^, 5/41 (12.2%) with BMI 25-30kg/m^2^, and 7/33 (21.2%) with BMI >30kg/m^2^ (21.2%), although this was not statistically significant (p=0.12).

**Table 1.** Demographics of the transplant cohort 2015. PRD: primary renal diagnosis; DBD: donation after brain death; DCD: donation after cardiac death, SCD: standard criteria donor, ECD (extended criteria donor).

| Summary demographics | Number of Patients (total 132) | Percent / Range |
| --- | --- | --- |
| **Recipient:** |  |  |
| Average age (years) | 49.9 | 19 to 74 |
| Male | 79 | 59.8% |
| **Body Mass Index (BMI)#** |  |  |
| BMI $<25\text{kg/m}^2$ | 58 | 43.9% |
| BMI 25-30$\text{kg/m}^2$ | 41 | 31.1% |
| BMI $>30\text{kg/m}^2$ | 33 | 25% |
| Diabetes as PRD | 16 | 12.1% |
| Live Donation | 40 | 30.3% |
| Deceased donation: | 92 | 69.7% |
| DBD | 58 | 63.0% |
| DCD | 34 | 36.9% |
| SCD | 62 | 67.4% |
| ECD | 30 | 32.6% |
| Average donor age (years) | 51.3 | 11 to 77 |

### Clinical contact days

Inpatient days, outpatient clinics, and day ward attendances accounted for 7035 episodes during the 2-year observation period; each of these episodes is visually represented in Figure 1. The median number of clinical contact days per patient was 47 (with medians of 12 in-patient, 3 day ward, and 28 out-patient appointments). Clinical contact days over the 2 years of follow up were front-loaded, with a total of 3235/7035 (46.0%) occurring in the first 90 days post-transplant, see Figure 1. Figure 1 also demonstrates the variability between patients, with 16/132 individuals (12.1%) experiencing less than 30 clinical contact days, compared to 13/132 (9.8%) with greater than 90 clinical contact days; the variation arising from the in-patient days (IQR 15.8 days), with fairly homogenous day ward and outpatient activity – see Figure 1 and Table 2. Identifying the characteristics of this cohort with heavy bed usage, both BMI >30kg/m^2^ and ECD subgroups contribute: BMI >30kg/m^2^ median 15.5 in patient days, versus 12.0 with BMI <25kg/m^2^, p=0.05 (data were also skewed, with mean in-patient days of 27.9 and 17.0 respectively); ECD subgroup accrued median 18.0 in-patient days versus SCD 11.0 days, p<0.001. The differences in subgroups could even be demonstrated at the level of length of post-operative admission, median days being 7 for both SCD and live donation, and 10 for ECD (P=0.001); and 7 days with BMI <25kg/m^2^, 8 with BMI 25-30kg/m^2^, and 9 if >30kg/m^2^, P=0.02. DBD and DCD versus live donation did not influence admission days (deceased donor median 12 days, live donor 14 days). Neither day ward nor out-patient attendances differed by subgroup (see Table 2).

**Figure 1.**
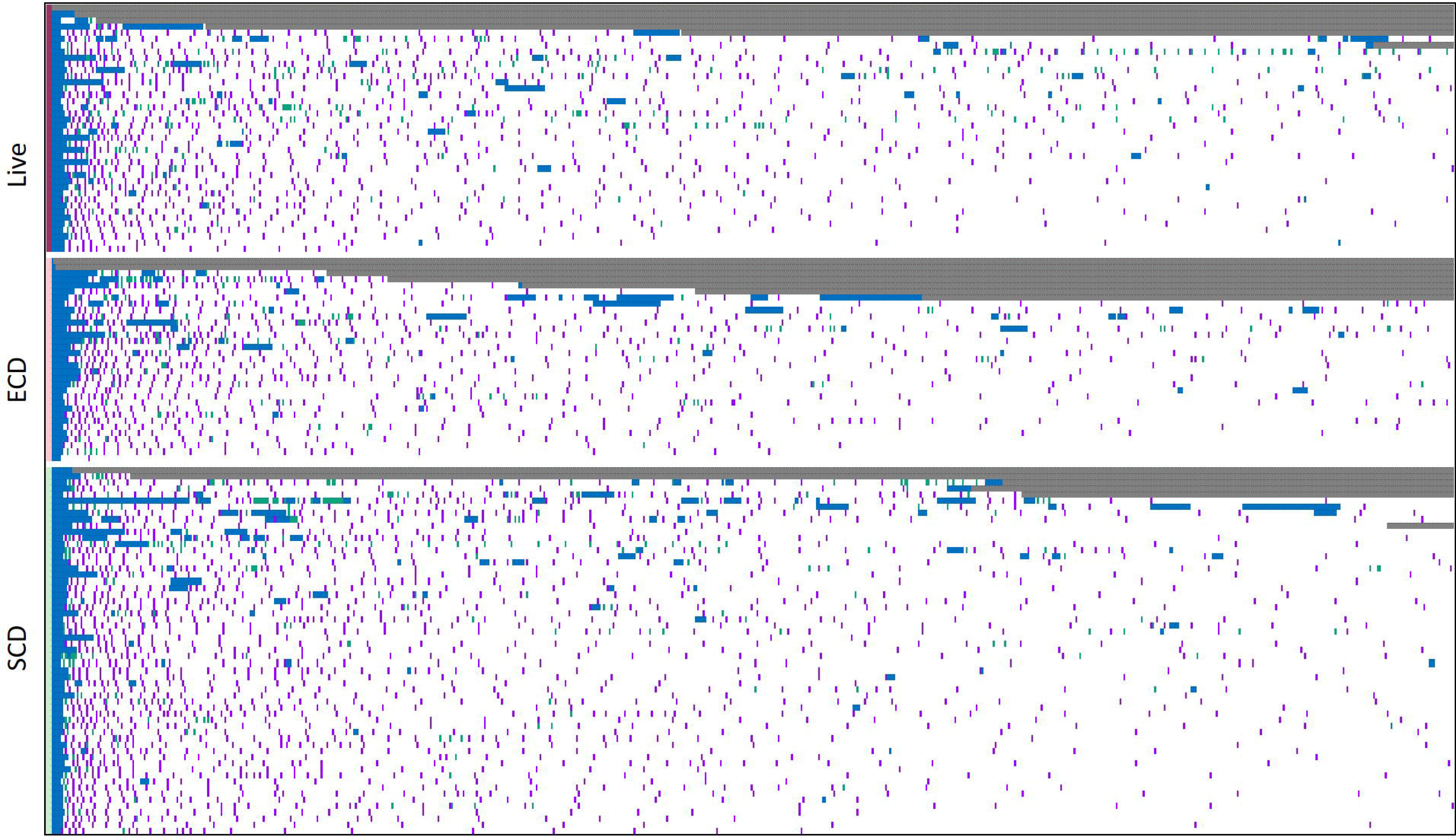
Clinical contact days by patient (rows) and day number (columns), sub-grouped by Live, Standard criteria donation (SCD) and Extended criteria donation (ECO). Blue: in-pat ient day; Purple: out-patient attendance; Green: day ward attendance; Grey: censored (death or graft loss).

**Table 2.**
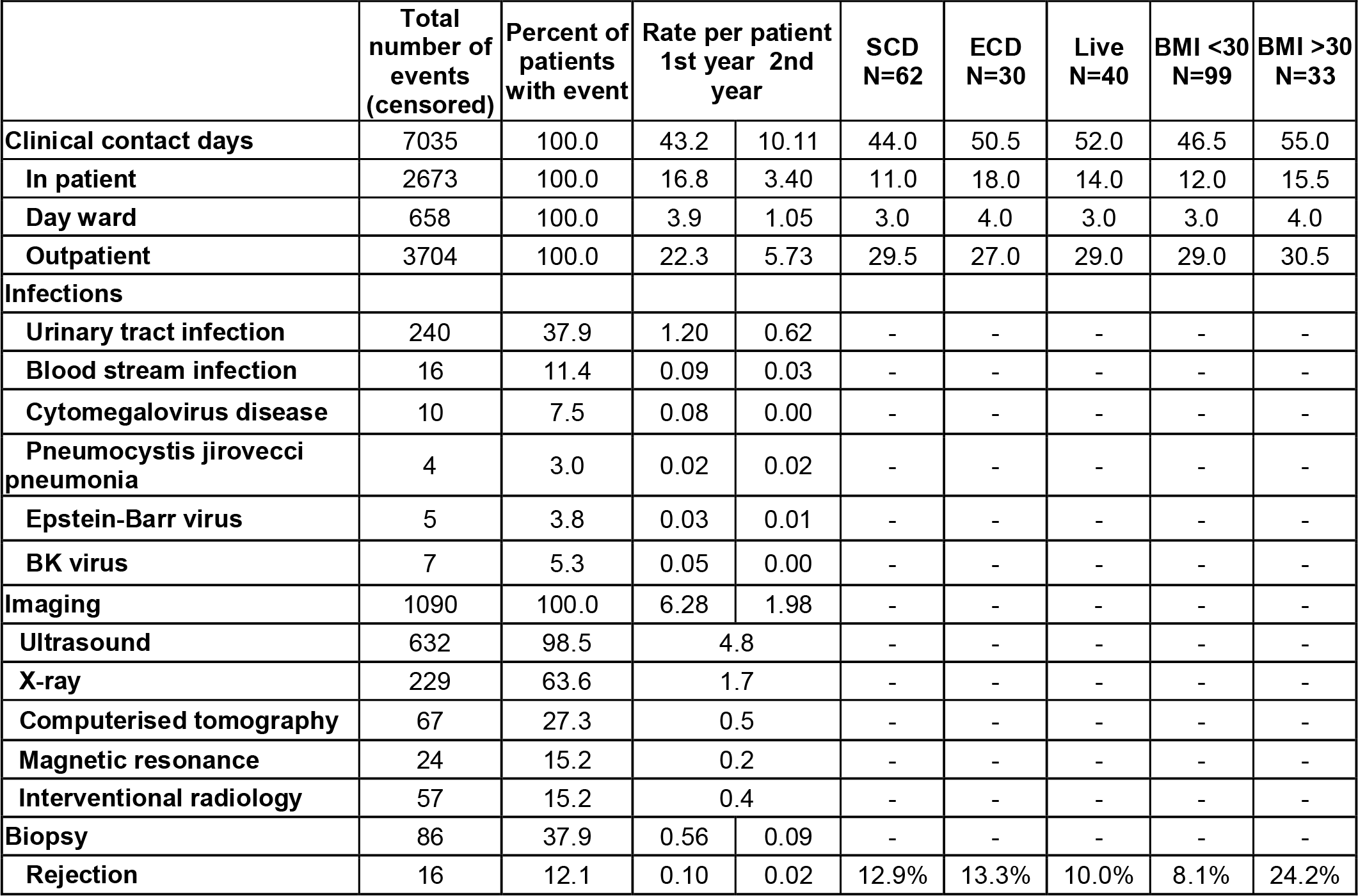
Healthcare activity events and clinical contact days over 2 years following transplantation.

### Graft loss

Over the 2 years, 11/132 (8.3%) individuals lost graft function and returned to dialysis, and 8/132 (6.1%) recipients died. One-year patient survival was 128/132 (97.0%). One-year graft survival was 123/128 (96.0%). Recipients of ECD kidneys had higher rates of graft loss and death in the first year (combined outcome 10/30 (33.3%) versus 2/62 SCD (3.2%), HR 10.3 (95% CI 2.4 – 44.2) P<0.001, compared with 1/40 (2.5%) of live donor transplants. However, in the second year the divergence disappears (0/20 remaining ECD kidneys, versus 3/60 SCD (5%), and 3/39 live donor (7.7%). Considering kidneys as live donation, DBD, or DCD, the combined outcome of graft failure or death doubled, from 5/40 (9%) of live donation, to 8/58 (13.8%) of DBD kidneys, to 6/34 (17.7%) of DCD kidneys (NS, p=0.93).

### Infection

64/132 (48.5%) of the cohort experienced at least one episode of microbiological or virologically proven clinically significant infection, whilst 9/132 (6.8%) experienced 10 or more distinct infection episodes. 50/132 (37.9%) experienced at least one confirmed UTI, the temporal distribution of infections is illustrated by Figure 2. 16/132 (11.4%) developed a blood stream infection. 4/132 (3.0%) developed PJP; all after the standard 6 month course of prophylactic co-trimoxazole had been discontinued. 10/132 (7.6%) developed CMV disease with all cases occurring in the first year post transplant, of which 3 patients were on prophylactic valganciclovir at the time of diagnosis. EBV infection was diagnosed in 5/132 (3.8%), and BK infection in 7/132 (5.3%). No difference in infection rates was detected by donation type (live versus cadaveric, or ECD versus SCD).

**Figure 3.**
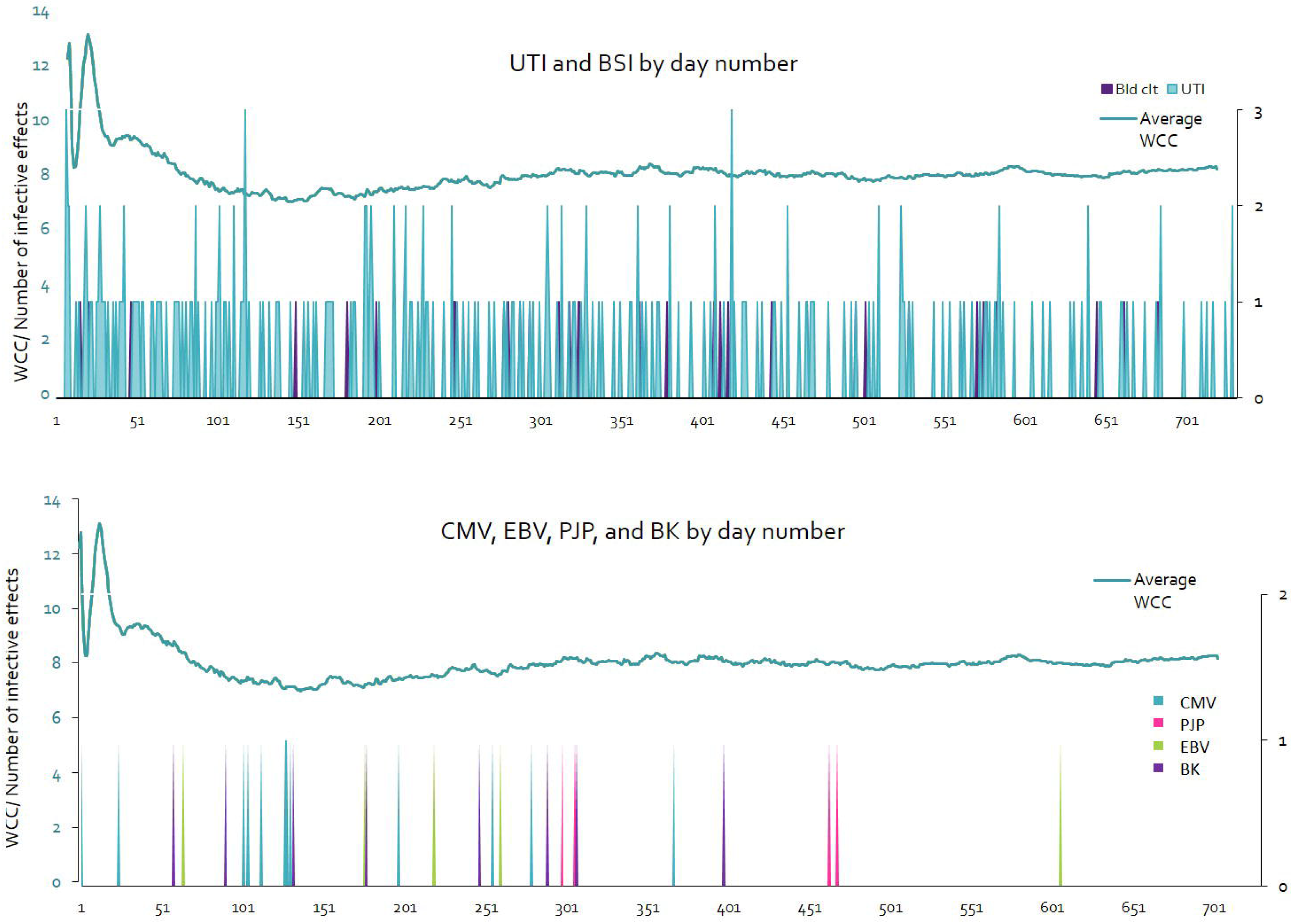
Infection complications. Events by day number. UTI: urinary tract infection; BSI: blood stream infection; WCC: White cell count; Bid cit: blood culture; e MV: cytomegalovirus-relateddisease; PJP: pneumocystis jiroveccipneumonia, EBV: Epstein Barr virus related-disease, BK: BK nephropathy

Infections were weighted to the first year (68.4% of events), as illustrated in Figure 2, where the day-by-day average patient WCC is also illustrated. Days 141-153 post-transplant corresponded to a relative nadir in WCC of less than 6.6×10^9^/L. 61/132 patients (46.2%) had an episode of leukopaenia (WCC <4×10^9^/L) during two years of follow up.

### Imaging

A total of 1090 completed imaging studies were undertaken over the 2 year observation period following transplantation; broken down as 632 ultrasound studies (57.9% of the 1090 imaging events), 229 X-rays (21.0%), 67 computerised tomography (CT) scans (6.1%), 57 interventional radiology procedures (5.2%), 24 magnetic resonance studies (2.2%), and 81 miscellaneous or uncategorised imaging events (e.g. nuclear medicine studies, video fluoroscopy, dual-energy X-ray absorptiometry, isotope bone scan etc.). 540/1090 (49.5%) of imaging studies occurred in the first 90 days.

### Biopsy and Rejection

86 kidney transplant biopsies were undertaken in 50/132 recipients (37.9%) over the 2 years. 45/86 (52.3%) of the biopsies occurred in the first 90 days. Of these, 5/45 (11.1%) showed rejection compared to 17/41 (41.4%) of those biopsies undertaken after 90 days. Overall a biopsy proven rejection episode occurred in 16/132 (12.1%) of patients, two antibody-mediated, who received optimised immunosuppression, and 14 with cell-mediated rejection, all of whom received methyl-prednisolone and two received anti-thymocyte globulin (ATG). ECD kidneys had similar rates of biopsy to SCD, and similar rates of rejection: 8/62 (12.9%) of SCD recipients versus 4/30 (13.3%) of ECD recipients). DCD kidneys were more likely to be biopsied, with 18/34 (52.9%) undergoing biopsy compared to 22/58 (37.9%) of DBD and 10/40 (25%) of LD kidneys (p=0.05), but with no difference detected in rejection rates by live versus deceased donation type, or by BMI.

### PTDM

12/132 recipients (9.1%) developed PTDM, after a median 174 days. There was an excess of PTDM diagnosed in recipients with BMI >30kg/m^2^ (7/33 (21.2%), versus BMI <30kg/m^2^ 5/99 (5.1%); p=0.01).

## DISCUSSION

We report on a cohort of consecutive transplant patients from a service that serves half the Scottish population, with complete follow-up data across all related hospital, clinic, laboratory, and imaging services. Transplant services commonly collect patient and graft survival data for clinical audit purposes, though not data from these aforementioned domains which underwrite quality of care, but also a paradox of healthcare burden. Collating data across the many domains involved in post-transplant care is challenging; chronographic representations, such as Figures 1 and 2, can assist with portrayal of this multifaceted data. The cumulative effect of aggregating data in this way is to provide a granular map of what patients and their families may expect, and the areas of healthcare provision where greatest impact occurs. This is an important concept as it is imperative that patients should have a clear insight into the expected clinical journey following the initial transplant surgical procedure, and currently not all recipients do (15). Similarly, when considering healthcare provision in a time of expanding access to transplantation and ‘realistic medicine’, it is important for clinical governance that healthcare providers have an insight into all elements of down-stream healthcare interactions in the post-transplant period, and can configure services accordingly.

Whilst it is tempting to tether these details to financial data on cost, we have refrained from doing so for two reasons. Firstly, one would need to consider the whole transplant process, not just the post-transplant period; including attribution of workload - whether investigations were purposely sought as part of the transplant process or not (e.g. pre-operative coronary angiography, echocardiography); furthermore, healthcare activity undertaken in patients subsequently suspended or removed from the transplant waiting list should be considered. Secondly, our data are representative of the UK transplant population and model of care, internationally there exist differences both in patient demographics and structure of service provision (16–18), and so cost data may vary. By illustrating the activity per patient in each domain, the appropriate tariff within different healthcare economies may then be calculated. Finally, the true comparator to transplantation is dialysis; we have already mapped out one year of healthcare burden associated with an incident haemodialysis population (19). Despite immunosuppression, rates of bloodstream infection were lower in the first year of transplantation compared to a patients’ first year on haemodialysis; 0.09 per patient per year versus 1.19 (19). For incident patients commencing haemodialysis, the mean number of admission days in first year is 17.1 (19), compared to 16.8 days for transplant patients in their 1^st^ year, and 3.4 for the second year. Cost data on dialysis may vary in different healthcare economies.

## Post-Transplant Activity

With nearly 1 in 10 patients experiencing more than 90 clinical contact days in two years, and no clear predictors out with BMI and ECD status, patients should be prepared for the possibility of significant clinical contact with their transplant team; particularly as one of the main advantages over dialysis is the freedom transplantation usually provides. With each transplant incurring a median of 47 clinical contact days over 2 years, services that may experience an uplift in transplantation rates would require to have sufficient additional capacity across their inpatient, outpatient and day-area facilities to accommodate this.

Beyond the initial post-operative recovery period, a significant amount of clinical activity was spent dealing with the consequences of infection. The absolute rates of infections reported in this study were very similar to national registry data (20), and around half of our cohort experienced at least one clinically significant infection, two-thirds of infections occurring in the first year after transplantation. The biggest issue was that of urinary tract infection, with nearly 40% of patients experiencing at least one episode, and studies reporting associated increased risk of acute rejection (21); prevention, treatment and antibiotic stewardship of urinary tract infection following transplantation must therefore remain a clinical priority. A minimum of 1 in 12 patients appeared to experience significant viral infections, whilst PJP was a relatively rare but nonetheless major event with associated mortality risk.

On average 8.2 imaging studies were undertaken per patient, with much of this activity (57.9%) being USS, and activity weighted to the first 90 days (49.5% of all imaging studies undertaken over the two years). Exposures to imaging requiring ionising radiation episodes were frequent, with 1.7 X-rays and 0.5 CT scans per patient over the 2 years, as well as a broad range of interventional radiology, magnetic resonance studies, and other activities. Availability of a range of imaging modalities, both cross-sectional and interventional, appears to be heavily relied upon by transplant services.

Almost 40% of the cohort required renal biopsy, with half of those biopsies occurring within the first 90 days. This reflects a reactive, rather than protocolised, approach to post transplant biopsy in our service. The probabilities of identifying acute rejection appeared to change over time with a lower proportion of early post-transplant biopsies identifying rejection, however this is qualified by the significance of missing early rejection in a newly placed graft.

The excess of PTDM diagnosed in recipients with BMI >30kg/m^2^ (21.2% (7/33) versus 5.1% with BMI<30kg/m^2^), must be considered alongside the additional 7/33 who already had a primary renal diagnosis of diabetic nephropathy, and others who were recipients of simultaneous kidney pancreas not included in this analysis. Hence 42% (14/33) or more of kidney transplant recipients with BMI >30kg/m^2^ may have diabetes mellitus. Data from our Transplant centre between 1994 and 2004 found a rate of PTDM of 7.7% (22), compared to the overall rate here of 9.1%. This may be due to different diagnostic criteria, increased awareness and diagnosis, or a true rise in prevalence and requires further investigation.

## Applicability

The demographics, peri-transplant care, and outcome data for this cohort are reported alongside the activity data to allow assessment of their applicability to other populations. Specifically, rates of deceased versus live donation were very similar to UK-wide registry data (23,24). Similarly our rates of DCD and ECD kidney use were comparable with other transplant centres (20,23). Our finding of inferior graft and patient survival among ECD recipients in our cohort replicates that of other groups (25). Our study was intended however to illustrate the breadth of activity, rather than analyse intently the outcomes associated with donation categories, and key immunological and physiological influences on outcome are not taken into consideration, namely donor-specific antibodies and cold ischaemic times. Patients and service providers may be reassured that although ECD transplantation appeared to carry additional risk of complications and graft loss in the first year (and particularly in the first 90 days), ECD kidneys surviving beyond this period matched their SCD comparators in the subsequent year in our cohort.

Another important factor to consider is recipient BMI. Acceptance for transplantation in those with BMI greater than 30kg/m^2^ varies between transplant centers. In this cohort there was an excess of healthcare related activity associated with a BMI >30kg/m^2^, including additional in-patient days (mean 11 additional days in the first year post-transplant) and PTDM, but combined outcome of graft and patient survival did not differ by BMI category. Other studies do suggest higher risk of graft loss and patient death (12), though as we have demonstrated, there is a crossover of higher BMI and diabetes as the primary cause of renal failure, and teasing apart the relative contributions of each to the various outcomes is challenging.

Transplant kidney biopsy rates were comparable with national registry datasets, even when broken down by donor type. National rates of biopsy-proven acute rejection in the first year are quoted as 10.5% in Scottish registry data; this cohort demonstrated 10.6% in the first year (n=14), and 12.1% with acute rejection over 2 years (n=16). In this cohort, one year graft survival was comparable to the local and national averages (96.0%) (20,23).

In regards to the limitations, we are unable to compare subgroup differences in all activity domains, as for rarer events such as PJP, BK, and interventional radiology, the study would have been underpowered to do so. A few key outcomes measures with both patient and service implications are also notably absent from the study, particularly post-transplant malignancy and quality of life measures. Reported activity was also restricted to the post-transplant period, though pre-transplantation assessment and work up is also relevant for service planning and patient expectations but was not included here. Similarly, expanding the study to evaluate the healthcare activity associated with longer duration of transplantation, or any long term differences in the outcomes of ECD kidney transplant would also be worthwhile. Finally, as the field of transplantation continues to evolve, this study can only represent the current era, though in doing so can help inform service provision to improve quality of care, and to plan for changing demands.

## Summary

This study has mapped out the main clinical activity that occurs across a range of hospital inpatient, outpatient and procedural domains in the first two years following kidney transplant. Recipients, families, transplant, and radiology services should be geared to expect this high level of activity, influenced by recipient BMI and donation criteria, but reassured that activity falls beyond the first 90 days following transplant surgery.

## Patient consent for publication

not required.

